# RhoC modulates metabolic networks in cervical cancer by transcriptionally regulating the expression of genes involved in metabolism

**DOI:** 10.1101/2019.12.29.890640

**Authors:** Pavana Thomas, Chandra Bhavani, Sweta Srivastava

## Abstract

In their quest for autonomy, tumor cells are known to reroute metabolic networks to aid their proliferation and survival. These metabolic alterations are governed by the tumor sub-population, thereby contributing towards an additional layer of complexity within the already heterogeneous tumor. For instance, bulk proliferative tumor cells rely on completely different pathways for their metabolic requirements as opposed to the stem-like metastatic cells. However, the molecular switch that drives these metabolic changes remains unknown. RhoC is a well-established contributor towards multiple aspects of tumor development including proliferation, EMT, migration, invasion and metastasis. A transcriptomics-based approach on a RhoC overexpressing cervical cancer cell line unveiled distinct metabolic signatures existent in these cells. Oxidative phosphorylation, TCA cycle, nucleic acid metabolism and fatty acid elongation were some of the specific pathways that emerged as up-regulated. This study therefore provides insight into the intricate metabolic circuitry functional in aggressive RhoC-high cells and thus proposes a pivotal role for RhoC in oncometabolism.

## INTRODUCTION

The process of tumor progression and dissemination requires a network of events which regulate not just the phenotype specific to the stage of the disease, but also the molecular mechanisms which metabolically support the survival of cells. A cancer cell has a ‘mind of its own’ since it is blatantly disruptive, upsetting the body’s homeostasis at its worst and becoming autonomous at its best. In order to maintain its oncogenic addiction, it diverts all the channels of nourishment towards itself ensuring the ‘survival of the fittest’. The metabolic pathway is one of the means used by a cancer cell to its own end. Perturbations of metabolic pathways and formation of oncometabolites in cancer is long known. Genetic aberrations responsible for tumorigenesis can also affect metabolic pathways which can lead to accumulation of specific metabolites in the cancer cell, having enabling effects like increased survival. Tumor formation increases the diversity of metabolic networks depending on cell types. Be it the hypoxic cells at the core of the tumor, the cells facing challenges in the blood stream, or other cells at several stages of tumor progression, they are continuously facing a sea of changes in terms of the microenvironment. These cells are thus metabolically different and will have signaling machinery to support them. Thus, understanding the metabolic profile of tumor cell subsets would aid in developing modes to target these cells individually to maximize treatment efficacy. However, it is also important to note that presently there is minimal information on the metabolic distinctions between tumor cells and normal tissues.

Our work on the role of RhoC in tumor progression along with other abundant literature, support the contribution of RhoC in determining the sequence of events during tumor progression. Beginning from tumor formation till metastasis, RhoC is required for all the intervening steps, which include invasion, anoikis resistance, intravasation, extravasation and homing (Reymond et al., 2015; Srivastava et al., 2010; Thomas, Pranatharthi, Ross, & Srivastava, 2019). We have recently observed that RhoC provides radioprotection to tumor cells, a hallmark of cancer stem cells (Pranatharthi et al., 2019). The regulation of migration and invasion by RhoC has been shown in several tumors including breast, cervix, inflammatory breast cancer, lung and ovarian to name a few (Bravo-Cordero, Hodgson, & Condeelis, 2014; Chen, Cheng, Zhang, Li, & Geng,; Faried et al., 2006; Gou et al.,; Han et al., 2014; Li et al., 2013; Srivastava et al., 2010; Tumur et al.,; Vega, Fruhwirth, Ng, & Ridley, 2011; Yang et al.). Similarly the regulation of metastasis by RhoC has been well elaborated in several tumors (Clark, Golub, Lander, & Hynes, 2000; Hakem et al., 2005; Hoshino, Koshikawa, & Seiki, 2010; Iiizumi et al., 2008; Islam, Sharma, Kumar, & Teknos, 2013; Kleer et al., 2006; Kleer et al., 2002; Rosenthal et al., ; Wu, Wu, Kumar-Sinha, Chinnaiyan, & Merajver, 2004; Xing et al., ; Yao, Dashner, van Golen, & van Golen, 2006), where they show that alteration in the expression of RhoC results in changes in metastasis. Similarly, its role in regulating angiogenesis, anoikis resistance and cancer stem cell maintenance is undisputable (Thomas et al., 2019). The role of Notch1 in regulating RhoC in cervical cancer illustrates its upstream regulatory mechanism (Srivastava et al., 2010) as there is no driver mutation observed in RhoC as per the TCGA data, 2019. The regulation of MAPK, PYK and FAK has also been demonstrated as potential signaling networks instrumental in delivering the outcomes of RhoC signaling (Iiizumi et al., 2008). With such a vast and diverse functional consequence, we attempted to understand RhoC’s role in oncometabolism. Our present study affirmatively proposes its role in oncometabolism.

## MATERIALS AND METHODS

### Cell lines and culturing conditions

SiHa, a cervical squamous cell carcinoma (SCC) cell line was used in this study. Stable cell lines over-expressing RhoC (SiHa-R) and those containing the backbone vector alone (SiHa-N) were created. The cells were cultured using Dulbecco’s Modified Eagle’s Medium (DMEM) supplemented with 10% FBS (Fetal Bovine Serum) and maintained at 37°C in 5%CO_2_ conditions.

### Isolation of RNA

RNA was isolated using the TRIzol method as per the manufacturer’s protocol.

### Transcriptomic study

Illumina paired end sequencing (150×2) was used for the transcriptomic study. An average of 20.61 million Illumina HiSeq high quality reads were obtained, with 91.77% aligning to the human reference genome. Tophat-2.0.13 was used for reference alignment and assembly and differential expression calculation was performed using Cufflinks-2.2.1. Cuffdiff was then utilized to identify significant changes in transcript expression. The transcriptomic analysis was performed in replicates of n=2.

### Bioinformatic Analysis

DAVID (The Database for Annotation, Visualization and Integrated Discovery, version 6.8) was used for gene ontology and pathway analysis for the significantly up-regulated genes in SiHa-R cells. The heatmap was generated using Clustvis, an R-based bioinformatic tool. Network analysis was performed using the STRING database.

## RESULTS

### Transcriptomic analysis reveals that SiHa-R cells exhibit pronounced metabolic signatures

Tumor cells are known to modify and adapt themselves facilitating growth, dissemination, invasion, migration or anoikis-resistance mainly determined by the stage of the tumor and the sub-population in question. Depending on the cellular phenotype, nutrients and metabolic resources are channelized either into rampant anabolism leading to rapid cell division, or limited metabolic activity barely sufficient for survival, as is the case with hypoxic cells. Intrinsic molecular pathways existent in these cells drive their metabolic activity. The protein RhoC has been shown to be involved in multiple tumor phenotypes in cervical cancer including that of proliferation, invasion, migration, anoikis-resistance and resistance to radiation therapy (Pranatharthi et al., 2019; Srivastava et al., 2010). Transcriptional profiling of SiHa cells (a cervical carcinoma cell line) over-expressing RhoC (SiHa-R) in comparison to SiHa cells containing the background vector alone (SiHa-N), indicates global up-regulation of genes in SiHa-R cells (Pranatharthi et al., 2019). Further analysis of this data divulges the significantly enriched metabolic status of SiHa-R cells.

KEGG pathway analysis available on DAVID (The Database for Annotation, Visualization and Integrated Discovery, version 6.8), was used to identify the pathways that were enriched by the significantly up-regulated genes in SiHa-R cells. As shown in **Figure 1a**, pathways involved in metabolism emerged as having the maximum number of genes in the up-regulated group (148 out of 1627). The other pathways that were enriched in SiHa-R cells were those involved in RNA transport, Huntington’s disease, Alzheimer’s disease, Parkinson’s disease, spliceosome, DNA repair and cell cycle among several others. GO analysis of the cellular component (CC) of the up-regulated genes in SiHa-R revealed localization of a significant number of these gene products in the mitochondria and endoplasmic reticulum, as illustrated in **Figure 1b**. A total of 220 proteins localized to various sub-organellar regions within the mitochondria (matrix, inner membrane, inter-membrane space and outer membrane) whereas 83 proteins were known to be functional within the endoplasmic reticulum. In order to understand the biological processes (BP) that the up-regulated genes in the metabolic category are involved in, the metabolic genes enriched by pathway analysis of the over-expressed genes in SiHa-R were reanalyzed using the BP tool available under GO analysis in DAVID. ATP synthesis coupled proton transport, redox processes, processes involved in the tricarboxylic acid (TCA) cycle, glycosylphosphatidylinositol (GPI) anchor biosynthesis (involved in directing proteins to the endoplasmic reticulum), ATP hydrolysis coupled proton transport and mitochondrial electron transport emerged as the key biological processes that were significantly increased in SiHa-R cells (**Figure 1c**). Molecular function (MF) analysis of the metabolic genes placed hydrogen ion transmembrane transporter activity, NADH dehydrogenase activity, proton-transporting ATPase activity, cytochrome-C oxidase activity, electron carrier activity and 5’-nucleotidase activity among the top molecular events that these genes regulate, further emphasizing the metabolically active status of SiHa-R cells (**Figure 1d**). Taken together, these data reiterate that there are specific metabolic pathways that are selectively activated in SiHa-R cells, pointing towards a possible role of RhoC in regulation of metabolism in cancer cells.

**Figure 1:**
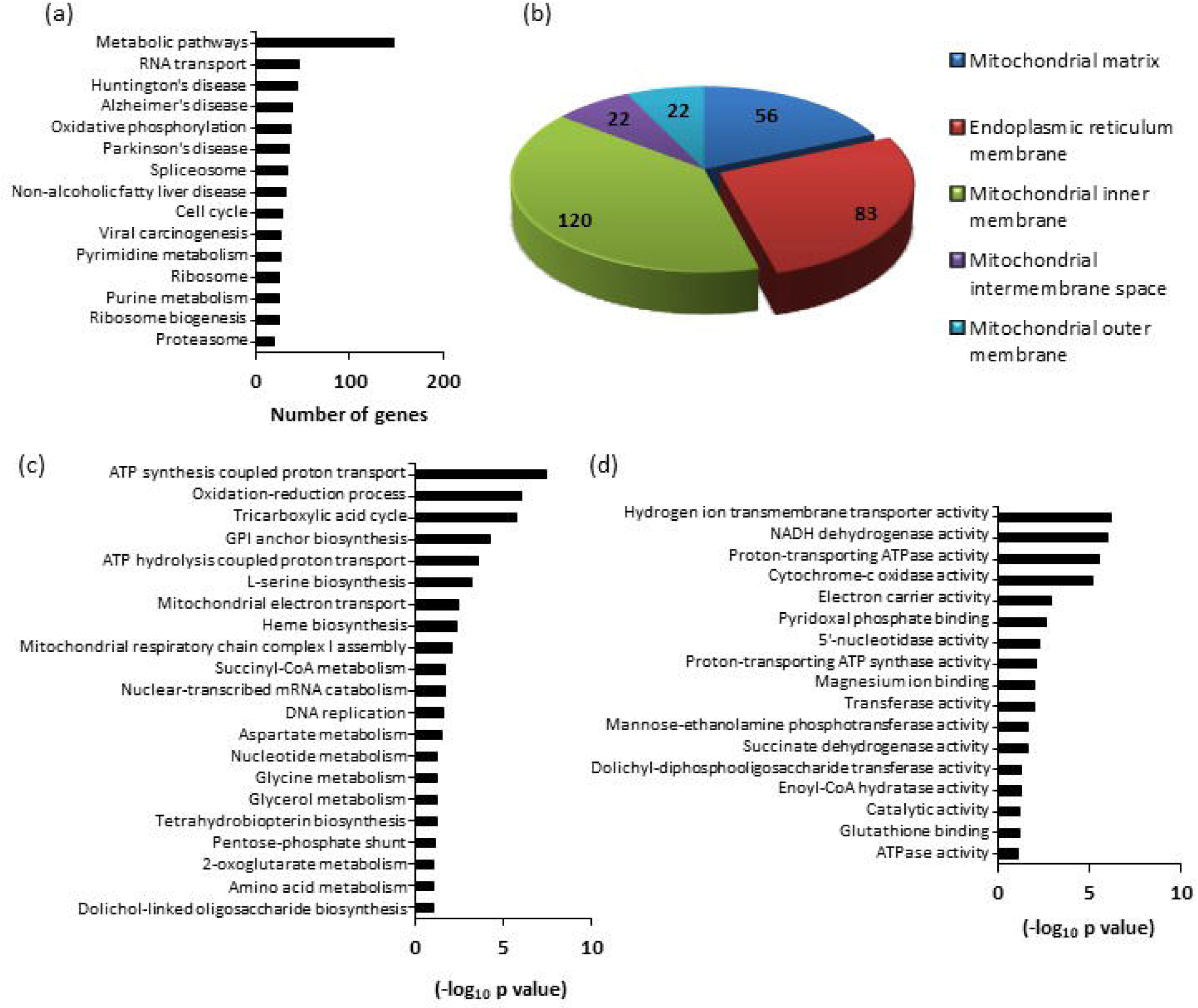
SiHa-R cells show significant up-regulation of metabolic pathways and processes-. (a) Bar graph representation of the top pathways functional in SiHa-R cells as revealed by KEGG pathway analysis of genes significantly up-regulated in SiHa-R cells. The maximum number of genes were found to be involved in metabolic pathways, indicating a prominent role for RhoC in metabolism. (b) A pie chart showing the cytosolic distribution of the genes involved in metabolism in SiHa-R, with a large majority of them localizing to the mitochondria and the endoplasmic reticulum. (c) Graphical representation of the prominent biological processes regulated by the up-regulated gene set in SiHa-R. (d) Graphical representation of the major molecular functions of the up-regulated genes in SiHa-R.

### Enrichment of genes associated with metabolic pathways in RhoC over-expressing cells

In order to determine the distinct pathways existent within cells, it becomes crucial to decipher the molecular interactions that occur between proteins. Towards this end, the 148 genes that were up-regulated in metabolic pathways were fed into the STRING database (Szklarczyk et al., 2015) to identify the pivotal functional protein interactions that take place among the metabolic proteins that were enriched in SiHa-R cells. It was found that the genes involved in metabolism which were over-expressed in SiHa-R cells distinctly clustered into the oxidative phosphorylation, TCA cycle, purine/pyrimidine metabolism, fatty acid elongation and carbon metabolism pathways (**Figure 2a**). This provided a coherent understanding of the specific molecular networks active in cells over-expressing the RhoC protein. The fold changes in gene expression of metabolic genes classified under distinct pathways was used to generate a heatmap using the web-based tool, Clustvis (Metsalu & Vilo, 2015). **Figure 2b** unambiguously illustrates the up-regulation of metabolic genes involved in oxidative phosphorylation, nucleic acid metabolism, carbon metabolism, N-glycan biosynthesis, glutathione metabolism and biosynthesis of unsaturated fatty acids in SiHa-R cells as compared to SiHa-N cells.

**Figure 2:**
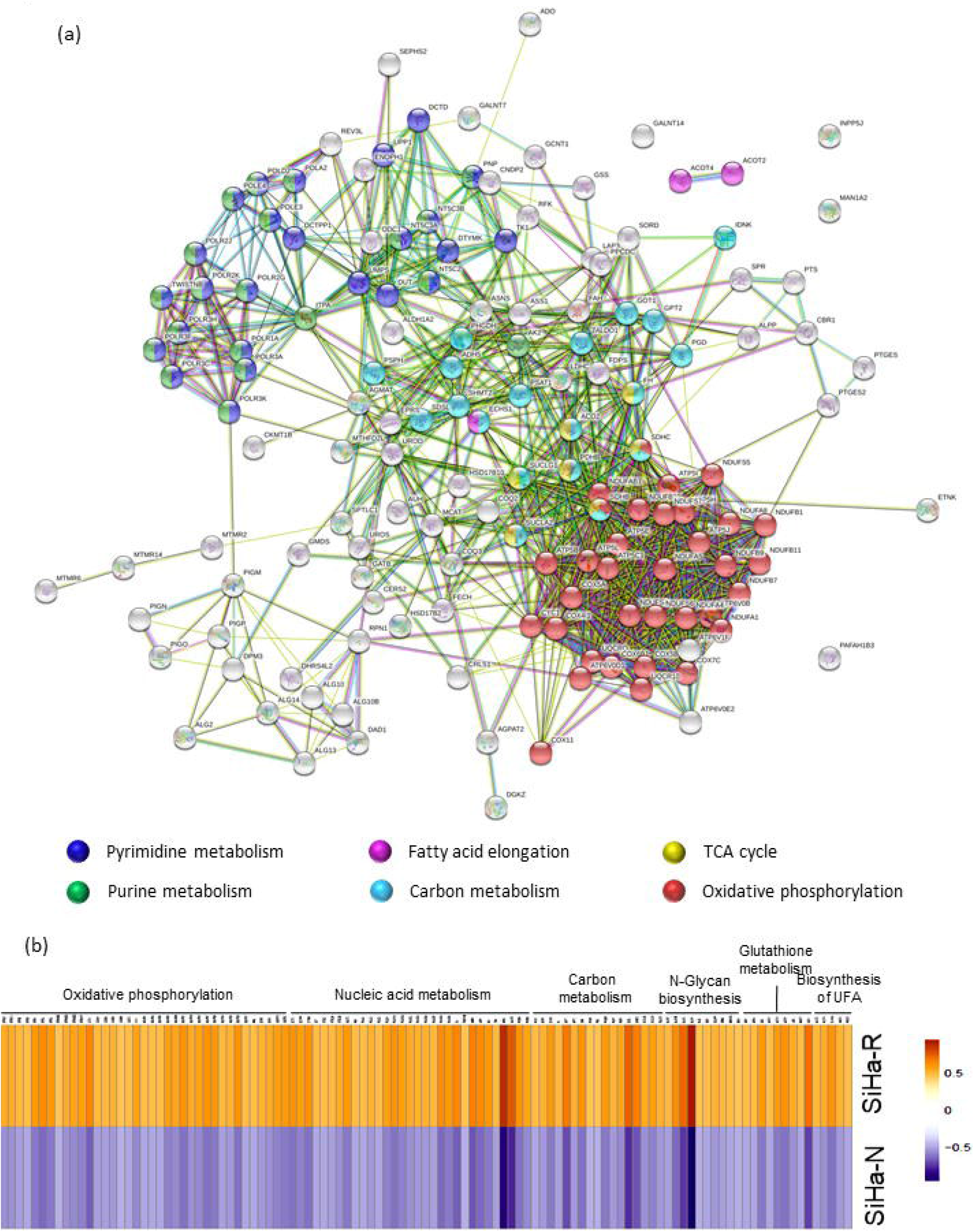
An overview of the metabolic pathways enriched in SiHa-R cells-. (a) STRING analysis of the up-regulated genes involved in metabolism with genes organizing themselves into discrete clusters based on the pathways that they are involved in. (b) Heatmap representation of FPKM values of genes involved in oxidative phosphorylation, nucleic acid metabolism, carbon metabolism, N-glycan biosynthesis, glutathione metabolism and biosynthesis of unsaturated fatty acids (UFA).

### Oxidative phosphorylation and TCA cycle pathway signatures are strongly up-regulated in SiHa-R cells

To identify precise alterations in the molecular machinery which allow hyperactivation of the oxidative phosphorylation pathway in SiHa-R cells, a molecular function based STRING network analysis was performed on the genes enriched in SiHa-R cells that are involved in oxidative phosphorylation. As shown in **Figure 3a**, proton transmembrane transporter activity, NADH dehydrogenase activity, cytochrome-c oxidase activity, co-factor binding, electron transfer activity and oxido-reductase activity were the major molecular processes that were regulated by these genes. The fold changes of these genes in SiHa-R are represented in **Figure 3b**. The NDUF gene family codes for the NADH:ubiquinone oxidoreductase proteins which serve as the entry point for electrons in the electron transport chain in mitochondria. This enzyme catalyzes removal of electrons from NADH and transfers them to the electron acceptor, ubiquinone (Walker, 1992). The genes ATP5B, ATP5E, ATP5H, ATP5I, ATP5J, ATP6V0B, ATP6V0D1 and ATP6V0E2 encode for sub-units of the ATP synthase protein which are responsible conversion of ADP to ATP utilizing energy generated from the electron transport chain. The Cox group of genes is translated to cytochrome-c oxidase proteins which function as the terminal electron acceptor in the electron transport chain. Cytochrome-c oxidase has been shown to be regulated by the Ras oncoprotein and is positively associated with development of poorly differentiated tumors in mice in lung adenocarcinomas (Telang et al., 2012). Succinate dehydrogenase B (SDHB) is a subunit of the enzyme complex involved in transferring electrons from FADH to CoQ, while succinate dehydrogenase C (SDHC) helps in anchoring the protein complex to the inner mitochondrial membrane (Cardaci & Ciriolo, 2012). UQCR10 and UQCRQ are ubiquinone binding proteins that are components of complex III of the mitochondrial electron transport chain (Usui, Yu, & Yu, 1990).

**Figure 3:**
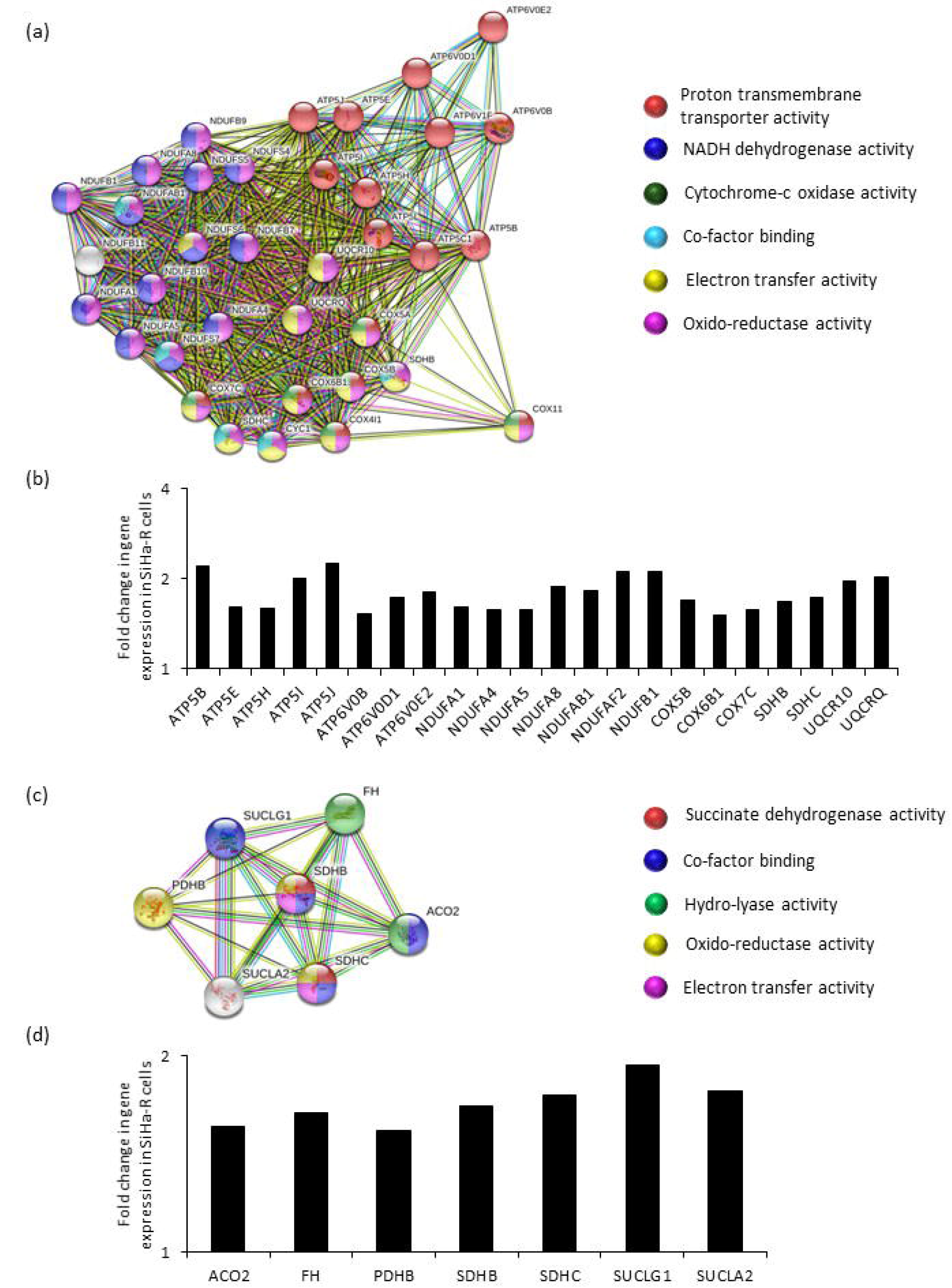
Analysis of molecular functions of genes involved in oxidative phosphorylation and TCA cycle-. (a) STRING analysis of up-regulated genes involved in oxidative phosphorylation reveal the major molecular processes that these genes govern. (b) Graphical representation of the fold changes of some of the oxidative phosphorylation genes enriched in SiHa-R. (c) STRING network analysis of the enriched genes that clustered in the TCA cycle depicting their prominent molecular functions. (d) Bar graph of the fold changes of some TCA cycle genes up-regulated in SiHa-R.

A similar STRING analysis of molecular functions on genes clustered under the TCA cycle illustrated that the enriched genes have succinate dehydrogenase activity, co-factor binding capability, hydro-lyase activity, oxido-reductase activity and electron transfer activity (**Figure 3c**). **Figure 3d** represents the fold changes in expression of these genes observed in SiHa-R cells. Fumarate hydratase or fumarase (FH) is a component of the TCA cycle which catalyzes the addition of a water molecule to fumarate, giving rise to malate (Scott & Powell, 1948). SUCLA2 and SUCLG1 code for different sub-units of the succinyl-CoA ligase enzyme which catalyzes the conversion of succinyl-CoA to succinate, leading to the generation of one ATP molecule in the process (Leitzmann, Wu, & Boyer, 1970). The PDHB gene product is the beta subunit of pyruvate dehydrogenase involved in the first step of the TCA cycle and allows conversion of pyruvate to acetyl-CoA, which further feeds into the citric acid cycle upon conversion to citrate (Coxon, Liebecq, & Peters, 1949). ACO2 codes for aconitase, an isomerase that converts citrate to iso-citrate via the intermediate cis-aconitate (Krebs & Holzach, 1952).

Overall, analysis of the transcriptome of SiHa-R cells reveal a significant up-regulation of genes involved in both oxidative phosphorylation and TCA cycle, which is in agreement with emerging literature which implicates oxidative phosphorylation to be the prime metabolic mechanism functional in the therapy-resistant, stem cell sub-population in tumors (Pasto et al., 2014; Porporato, Filigheddu, Pedro, Kroemer, & Galluzzi, 2018; Viale et al., 2014; Weinberg et al., 2010).

### SiHa-R cells have robust nucleic acid metabolism and fatty acid elongation/synthesis signatures

Rapid, deregulated cellular proliferation is a hallmark of cancer cells. This requires abundant and continuous supply of nucleotides which serve as building blocks for DNA replication and transcription of genes to RNA. The up-regulated genes in SiHa-R cells that are involved in purine/pyrimidine metabolism were further analysed for molecular functions and clustered using STRING. The analysis revealed that DNA-directed DNA polymerase activity, nucleotidyl transferase activity, DNA-directed RNA polymerase activity, nuleobase-containing compound kinase activity, 5’-nucleotidase activity and nucleoside monophosphate kinase activity were the key molecular processes that were regulated by these genes (**Figure 4a**). **Figure 4b** is a graphical representation of the increase in fold changes of these genes in SiHa-R cells in comparison to SiHa-N cells. Also, as represented by a venn-diagram (Venny 2.1.0), 19 genes that were overexpressed in SiHa-R cells were common to both purine and pyrimidine metabolism, whereas 7 genes were associated with pyrimidine metabolism only and 6 genes were involved in purine metabolism alone (**Figure 4c**). The Pol family of genes translates into enzymes that are involved in the extension of DNA/RNA strands (Billen, 1963; Furth, Hurwitz, & Anders, 1962). Consequently, all of them were seen to possess nucleotidyltransferase activity, while those involved in DNA and RNA polymerization formed two distinctly separate clusters as seen in **Figure 4a**. TWISTNB, another gene involved in DNA-directed 5’-3’ RNA polymerase activity was also found to have increased transcript levels in SiHa-R cells. The NT5 group of genes (5’ nucleotidases) are hydrolases that act on purines and catalyze their breakdown into a phosphate and the respective nucleoside, thus facilitating continuous recycling of the core molecules necessary for DNA/RNA synthesis (Heppel & Hilmore, 1951). TK1 (Thymidine kinase 1) is involved in the creation of deoxythymidine monophosphate (dTMP) from thymidine by the addition of a phosphate group, whereas DTYMK (deoxythymidylate kinase) converts dTMP to dTDP by the addition of another phosphate group (Eriksson, Munch-Petersen, Kierdaszuk, & Arner, 1991). AK2 (adenylate kinase 2) performs a similar function on adenosine by catalyzing the interconversion between ATP and AMP (Rosado et al., 1972).

**Figure 4:**
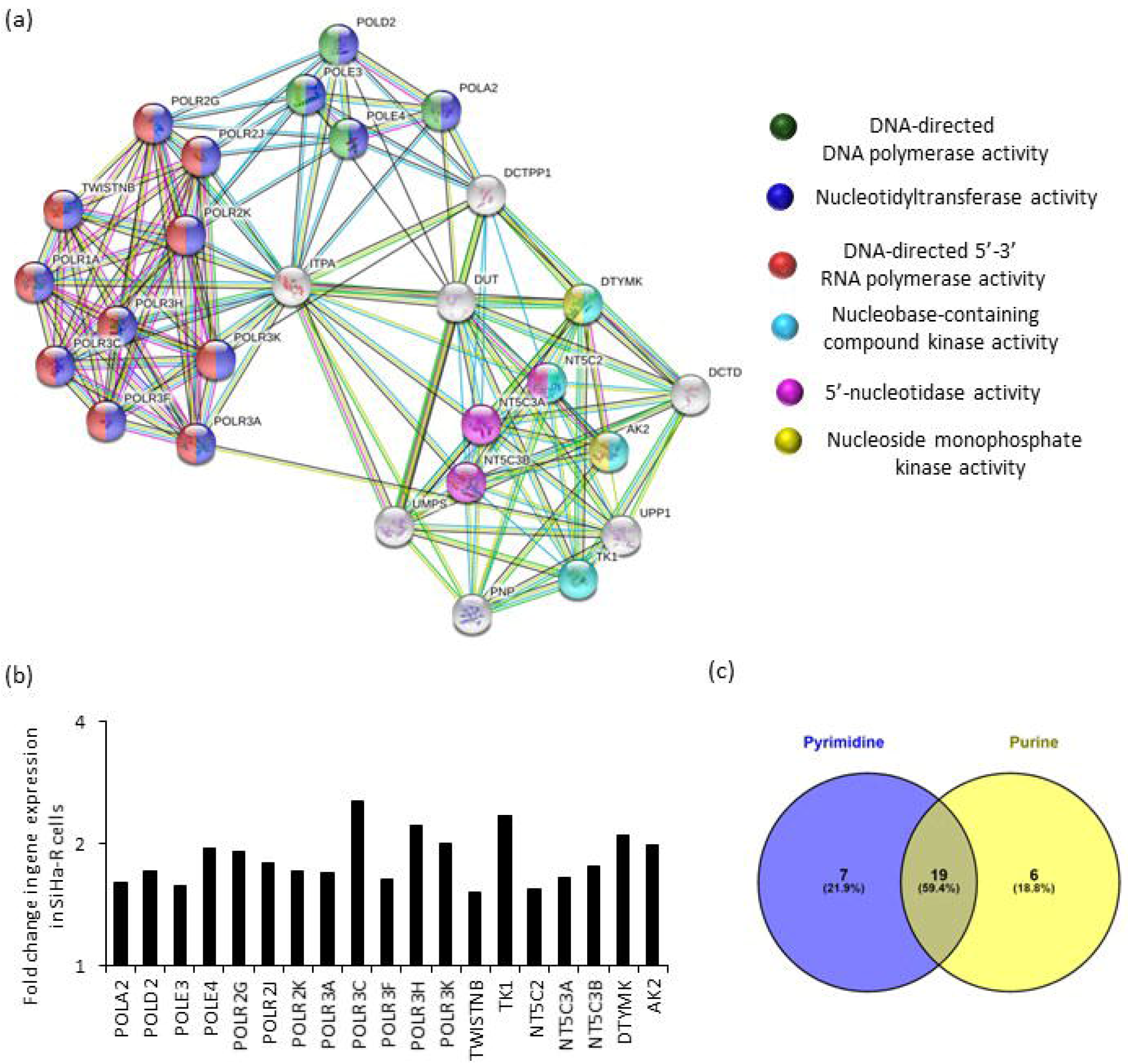
Evaluation of the molecular functions associated with genes regulating purine/pyrimidine metabolism-. (a) Network analysis using STRING depicts specific clusters of genes involved in purine/pyrimidine metabolism on the basis of their molecular functions. (b) Bar graph representation of the fold changes of selected genes in purine/pyrimidine metabolism that show increased levels in SiHa-R. (c) A venn diagram depicting the number of genes involved in purine metabolism alone, pyrimidine metabolism alone and common to both processes.

Proliferating cells require the synthesis of lipids as structural components for the plasma membrane, and membranes of other organelles within the cell. Apart from this, lipids also play a pertinent role in oncogenic signaling pathways. Processes involved in advanced stages of cancer like EMT (epithelial to mesenchymal transition), angiogenesis and evasion of the immune system depend on molecular signaling mechanisms that are largely regulated by lipids (Beloribi-Djefaflia, Vasseur, & Guillaumond, 2016; Kuo & Ann, 2018; Rohrig & Schulze, 2016). In fact, RhoC (also known to regulate invasion, migration and angiogenesis), requires prenylation for its membrane localization and activity (Zhang & Casey, 1996). Therefore lipid metaboilsm emerges as a critical factor in the realm of cancer biology. In our study, pathways involved in fatty acid metabolism were found to be significantly up-regulated in RhoC over-expressing SiHa cells. STRING analysis of the molecular functions of these genes indicated that they are associated with hydro-lyase activity, very long chain 3-hydroxyacyl-CoA dehydratase activity, fatty acid elongase activity, myrsitoyl-CoA hydrolase activity, palmitoyl-CoA activity and carboxylic ester hydrolase activity (**Figure 5a**). **Figure 5b** is a graphical representation of the increased fold changes of fatty acid elongation genes in SiHa-R cells. ECHS1 (Enoyl-CoA hydratase 1) is involved in conversion of enoyl-CoA to hydroxylacyl-CoA by the addition of a water molecule in the fatty acid beta oxidation pathway (Willadsen & Eggerer, 1975). PTPLA (also known as 3-Hydroxyacyl-CoA Dehydratase 1) and PTPLB (known as 3-Hydroxyacyl-CoA Dehydratase 2) are involved in synthesis of long chain and very long chain fatty acids. They catalyze the conversion of 3-hydroxyacyl-CoA to trans-2,3-enoyl-CoA by the removal of a water molecule, resulting in the addition of two carbon atoms to the fatty acid per cycle (Knoll, Bessoule, Sargueil, & Cassagne, 1999). ELOVL3 (Elongation Of Very Long Chain Fatty Acids 3) and ELOVL6 (Elongation Of Very Long Chain Fatty Acids 6) are also involved in fatty acid elongation and catalyze the addition of two carbon atoms from malonyl-CoA during one cycle of elongation (Jakobsson, Westerberg, & Jacobsson, 2006). ACOT2, ACOT4 and ACOT7 are acyl-CoA thioesterases which convert acyl-CoA to CoASH and the respective fatty acid, thereby regulating cellular levels of these molecules (Tillander, Alexson, & Cohen, 2017).

**Figure 5:**
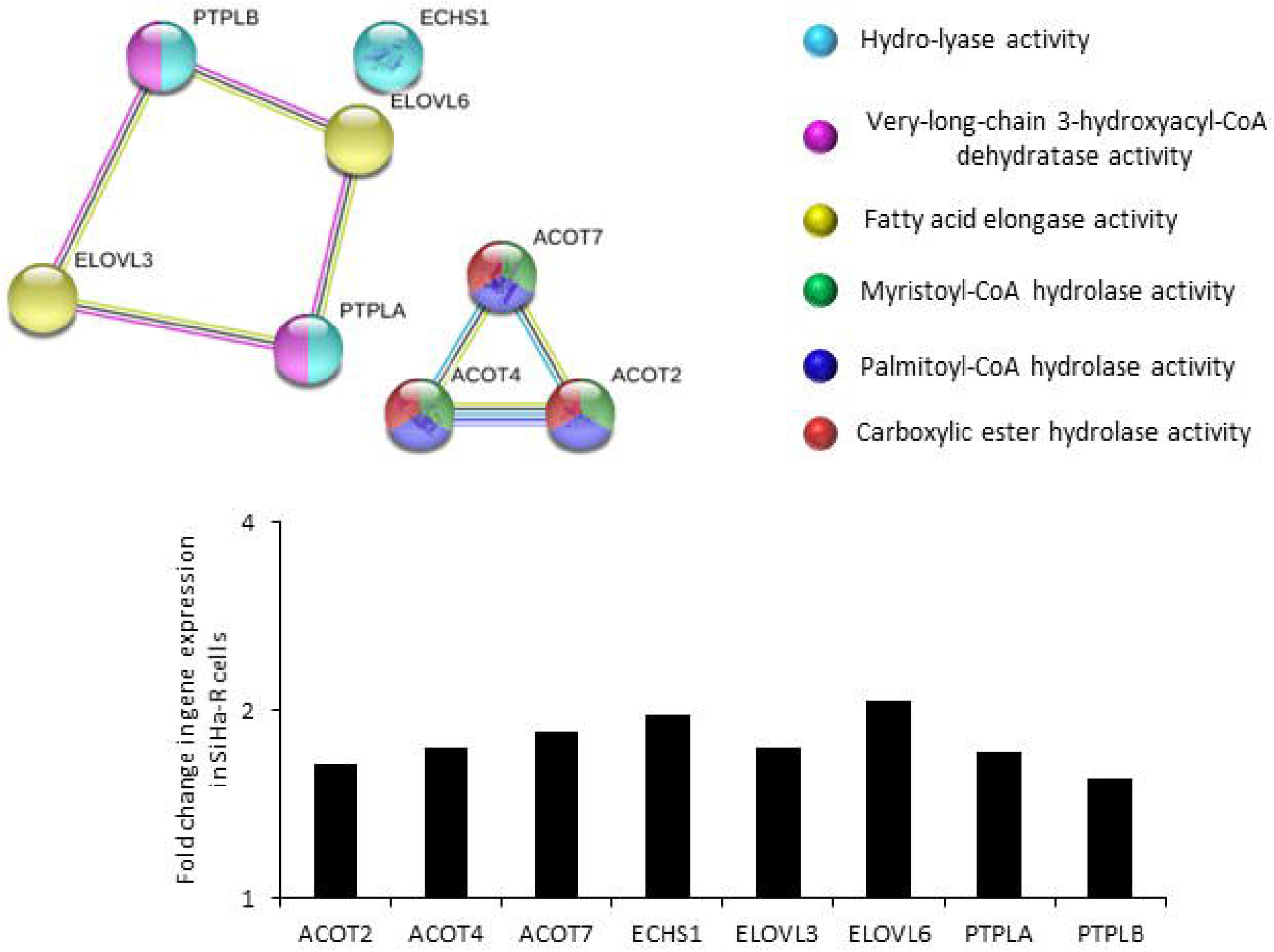
Investigation of the molecular activities of genes associated with fatty acid synthesis and elongation-. (a) Molecular function based STRING analysis illustrates the specific activities of genes up-regulated in SiHa-R cells that regulate fatty acid synthesis and elongation. (b) Graphical representation of the fold changes of genes associated with fatty acid synthesis and elongation in SiHa-R cells.

## DISCUSSION

The topic of cancer and its exploitation of the metabolic pathways came to the fore almost nine decades earlier (Morin, Letouze et al. 2014). In the 1930s when scientists tenaciously believed that only an external factor was responsible for cancer causation, Dr. Warburg believed otherwise. Dr. Warburg observed that proliferating mammalian cancerous cells in ascitic fluid utilized glucose even in the presence of abundant oxygen as the source of energy (Ward and Thompson 2012), which led to the formation of the eponymous hypothesis. Subsequently, mutations in genes involved in metabolic pathways like IDH1 (isocitrate dehydrogenase 1), IDH2 (isocitrate dehydrogenase 2) and SDH (succinate dehydrogenase) in cancer added weight to the hypothesis (Yang, Soga et al. 2012). The Warburg effect served as the rationale for the FDG-PET scan (^18^fluoro-2-deoxy-D-glucose (FDG) positron emission tomography (PET), an imaging technology which is used widely for detection and surveillance of tumors (Zheng 2012). Recent studies have shown that tumor heterogeneity reflects not only on phenotypes, but also has a major contribution to the realm of metabolic pathways that are active in these cells. Scientific evidence over the last few years corroborate that tumors do not always rely on a single metabolic process for their energy reserves. Additionally, data from multiple labs working on a range of tumor models have categorically exemplified that oxidative phosphorylation in the mitochondria is highly active and is associated with poor clinical outcome (Jones et al., 2016; Weinberg et al., 2010; Whitaker-Menezes et al., 2011). In fact, Viale A et al. have shown that pancreatic ductal adenocarcinoma (PDAC) cells that survive K-ras ablation, show stem-like properties and are highly dependent on oxidative phosphorylation for their metabolism (Viale et al., 2014). They further demonstrate that the metastatic cells (which arise from the cancer stem cell pool) are heavily reliant on oxidative phosphorylation and are therefore sensitive to inhibitors of the pathway, as opposed to the proliferative tumor bulk which largely use the glycolytic mode of metabolism.

RhoC is a protein that has been shown to be involved in varied tumor phenotypes including metastasis, EMT, anoikis resistance and radioresistance to name a few (Pranatharthi et al., 2019; Srivastava et al., 2010). To explore the role of RhoC in oncometabolism, a transcriptomic profile of cervical cancer cell lines-one over-expressing RhoC and its control, was performed. Based on the transcriptomic data, it was seen that RhoC regulates many processes like DNA doublestranded break repair and cell cycle progression (Pranatharthi et al., 2019).

Interestingly, of the 1627 differentially expressed genes, 148 genes were enriched for metabolic pathways. Some of the specified pathways which were up-regulated included oxidative phosphorylation pyrimidine metabolism, fatty acid elongation, carbon metabolism, purine metabolism, glutathione metabolism, citric acid cycle and biosynthesis of unsaturated fatty acids suggesting that the altered metabolic status of the cell is partially under RhoC regulation.

Our study revealed that genes involved in oxidative phosphorylation, spanning all the five mitochondrial complexes, were up-regulated. Studies have shown variable expression of genes belonging to Mitochondrial complex I (NADH:ubiquinone oxidoreductase) which is one of the largest proteins of the mitochondrial protein complexes which catalyze the formation of reactive oxygen species (ROS)(He, Zhou et al. 2013). Expression profile of NDUFS7, NDUFA1, NDUFA5 and NDUFAB1 in our study was comparable to other studies which demonstrated their over-expression in multiple myeloma cancer samples (Li, Sun et al. 2016; Mahajan, Wu et al. 2016), lung adenocarcinoma (Xu, Ma et al. 2016) samples, cervical cancer cell lines (Shimada, Moriuchi et al. 2006) and ovarian cancer samples (Verschoor, Verschoor et al. 2013) respectively. A bioinformatics revisit of the Oncomine cancer patient database shows an elevated expression of NDUFB1, NDUFB10, NDUFS4 and NDUFS5 in lung cancer patients (Li, Sun et al. 2016). NDUFS6 over-expression was observed in RNASeq data generated from breast cancer tissue samples, obtained from the Cancer Genome Atlas (Liu, Yin et al. 2017), comparable to the results of our study. However, contrary to what was observed in our study, a down regulation of NDUFA1 seems to be a pathogenic mechanism in basal cell carcinoma (Mamelak, Kowalski et al. 2005).

Similarly, the RhoC over-expressed cells exhibited an up-regulation of mitochondrial complex II (Succinate dehydrogenase/SDH) genes. An over-expression of SDHC was observed in BRCA-1 transfected breast cancer cell lines, which was indicative of activation of Krebs cycle and oxidative phosphorylation, (Privat, Radosevic-Robin et al. 2014) a result similar to that observed in the present study. Additionally, dysfunction of SDHB causes stabilization of hypoxia-inducible factor (HIF) which is responsible for angiogenesis in tumor cells (Xiao, Liu et al. 2018).

The present study also showed an elevated expression of CYC1 (Cytochrome c1) which belongs to Complex III of the electron transport chain called the cytochrome reductase or Q-Cytochrome C Oxidoreductase. Cytochromes are proteins which are a complex of heme and iron core (where iron can exist as the ferrous or the ferric form depending on the number of electrons it is carrying), and allows transfer of electrons from ubiquinol to cytochrome C which then transfers them to complex IV. Similar to what was observed in the present study, CYC1 of complex III was seen to be up-regulated in human osteosarcoma cell lines (Li, Fu et al. 2014), in the serum of osteosarcoma patients (Li, Zhang et al. 2009) and in breast cancer tissues of 3554 patients (Han, Sun et al. 2016). Our data shows a high expression of COX5B, a complex IV protein, which is in agreement with another study which concluded that an over-expression of COX5B (Cyclooxygenase 5B) was associated with poor prognosis in breast cancer patients (Gao, Sun et al. 2017). Increased gene expression of mitochondrial complex V in our data was also reflected in other studies which demonstrate an over-expression of ATP5B (adenosine triphosphate 5B) in ovarian cancer patients and ATPC1 (adenosine triphosphate C1) and ATP5I (adenosine triphosphate 5I) in breast cancer patients (Hjerpe, Egyhazi Brage et al. 2013, Sotgia, Whitaker-Menezes et al. 2012, Patsialou, Wang et al. 2012). Up-regulation of the mitochondrial gene expression in the RhoC over-expressed cells in our study, reiterates the hypothesis that RhoC is a positive and major metabolic facilitator in cancer cells.

Genes associated with another important metabolic pathway, namely Krebs cycle or the tricaboxylic acid cycle (TCA) was enriched in our data. Krebs cycle, existent in the mitochondrial matrix is involved in energy production by oxidation of acetyl-CoA. The Krebs cycle has many fuel sources like glucose, glutamine and fatty acids (Anderson, Mucka et al. 2018). It is very well documented that cancer cells have somewhat of a “sweet tooth” and depend heavily on glucose for their needs, thus shunting it away from the TCA. The end product of glycolysis is pyruvate which gets converted to lactate. The cancer cells are seen reusing and recycling lactate which gets reconverted to pyruvate and enters the Krebs cycle/TCA, thereby producing additional energy for the cancer cell (Bonuccelli, Tsirigos et al. 2010; Faubert, Li et al. 2017). The present study shows an over-expression of transcripts involved in TCA. The SDHB (succinate dehydrogenase complex iron sulfur subunit B) and SDHC (succinate dehydrogenase complex subunit C) transcripts also form the complex II of the mitochondrial respiratory chain and their involvement in tumorigenesis is mentioned above. ACO2 (aconitase 2) seen to be over-expressed in SiHa-R, is also up-regulated small cell tumor of lungs (Ocak, Friedman et al. 2014), whereas as it is under-expressed in prostate cancers (Tessem, Bertilsson et al. 2016). Contrary to our findings, FH (fumarate hydratase) down-regulation was observed in clear cell renal carcinomas (Sudarshan, Shanmugasundaram et al. 2011). Although the TCA genes have made their presence felt in many cancers, our study lends a unique dimension to this narrative by presenting RhoC as a protein that activates these metabolic pathways.

Additionally, the RhoC over-expressing cells show an enrichment of genes belonging to the purine/pyrimidine synthesis pathway, which maintains the nucleic acid pool aiding high cell turnover in neoplastic tissues. Up-regulation of various polymerases was observed in our study. High expression level of POLD2 (DNA polymerase delta 2, accessory subunit), which is a nucleotide excision repair gene correlated with poor overall survival and resistance to chemotherapeutic agents, in bladder urothelial carcinoma patients (Givechian, Garner et al. 2018). Elevated POLR2K (RNA Polymerase II Subunit K) expression levels in prostate cancer samples are associated with poor prognosis (Kelly, Sinnott et al. 2016). A survey of the Cancer Genome Atlas transcriptomic datasets show an up-regulation of TWISTNB (TWIST neighbour), a subunit of DNA-Directed RNA Polymerase Subunit I, in lung adenocarcinoma samples (Rossetti, Wierzbicki et al. 2016). Similarly, high gene expression levels of DCTD (deoxycytidylate deaminase) correlates with poor prognosis in malignant gliomas and could also be a potential therapeutic target in the same (Hu, Wang et al. 2017). DCTPP1 (dCTP pyrophosphatase 1) a Nucleoside triphosphate pyrophosphohydrolase (NTP) which maintains fidelity of DNA replication is seen to be up-regulated in our data. Similar overexpression of DCTPP1 in neoplastic cells has been observed in breast cancer cell lines (Zhang, Ye et al. 2013), gastric cancer cell lines (Xia, Tang et al. 2016) and is associated with tumor progression in prostatic cancers (Lu, Dong et al. 2018). Over-expression of the above genes involved in purine/pyrimidine metabolism in SiHa-R cells, help ensure that a constant supply of nucleotides is maintained to allow cellular proliferation and survival.

Gene expression of fatty acid biosynthesis and elongation showed an up-regulation in our study. Biosynthesis of unsaturated fatty acids and fatty acid elongation are two overlapping anabolic pathways. A cancerous cell acquires in its armatorium all the tools to make it self-sufficient, and one of them is de novo synthesis of fatty acids which is essential for the phospholipids. The otherwise healthy cells obtain their lipids from dietary or external sources (Currie, Schulze et al. 2013). We noted an over-expression of ELOVL6 (ELOVL Fatty Acid Elongase 6) in our study which is one the lipogenic markers that predicts poor prognosis, in terms of lymph node involvement and short recurrence free survival, in breast carcinoma patients (Feng, Chen et al. 2016). Further, its knock down in hepatocellular carcinoma cell lines inhibits cellular proliferation (Su, Feng et al. 2018). Similar to our data, ACOT2 (acyl-CoA thioesterase 2) expression, was increased in breast cancer cell lines with aggressive phenotype (Maloberti, Duarte et al. 2010) as was ACOT7 (acyl-CoA thioesterase 7), in human breast cancer and lung carcinoma cell lines (Jung, Lee et al. 2017). ECHS1 (enoyl-CoA hydratase, short chain 1), involved in beta oxidation of fatty acids was seen to be up-regulated in RhoC over-expressing cells. This molecule is capable of discerning the amount of nutrients, and accordingly allow activation of the mTOR (anabolic) pathway or down-regulation of the apoptotic pathway (Zhang, Qu et al. 2017). Consequently, an ECHS1 knockdown can have an inhibitory effect on cellular proliferation in gastric cancer cell lines (Zhu et al., 2014). Although the above mentioned genes have an increased expression in cancers, their notable up-regulation in RhoC over-expressed cancer cells forms a significant subtext of the entire document.

In the power play between the normal and neoplastic cells, most of the survival, anti-apoptotic and anabolic pathways are diverted to the cancerous cell in its pursuit of autonomy. The metabolic pathways too are utilized by cancer to facilitate its own growth and replenish its reserves. Dr. Warburg’s impressive scientific prescience led to more investigations in the area of cancer cell metabolism. As a result we have well-founded scientific documentation where scientists have shown that cancer cells utilize the fermentative pathways as surmised by Dr. Warburg along with utilization of the oxidative pathways, as shown in several studies as well as our own, that lends a new dimension to the entire narrative. The cancer cells are shown to be adept at upcycling their own waste and therefore the end products of metabolism are routed towards the TCA cycle which it uses to its advantage, as demonstrated in the present study. An addiction to the anabolic pathways is a straightforward logic which needs no justification. The fatty acid biosynthesis pathway and the pyrimidine metabolism pathways, both anabolic, are hyperactivated to compensate for the increased cell turnover, observed in the present study. A dysregulated expression of an oncometabolite can serve as a diagnostic marker, while a marker associated with the behavior of the tumor can be a potential prognostic marker. Interestingly this the first study which establishes that RhoC regulates these pathways by regulating the expression of the genes involved in these processes. This study emphatically shows that RhoC regulates cancer metabolism by regulating the gene expression. As represented by our data and several other groups, a cancer cell has numerous aberrant metabolic pathways. Thus developing therapeutics targeting every possible metabolic process is not viable option. Instead it would be meaningful to target a central signaling pathway which regulates all these processes. Our study reveals that RhoC is core to the regulation of these processes and inhibiting RhoC would be helpful in better clinical outcome.

## Acknowledgements

The transcriptomic study was funded by the Early Career Research Award awarded to Dr. Sweta Srivastava by the Science and Engineering Research Board, Department of Science and Technology, India. Travel grants from the Department of Science and Technology, India (DST-ITS) and Indian Council of Medical Research (ICMR), India have been awarded to Dr. Sweta Srivastava. Pavana Thomas has been awarded a fellowship from CSIR for the duration of her Ph.D.

## Data Availability Statement

Data is available on request from the authors. The data that support the findings of this study are available from the corresponding author upon reasonable request.

